# Experimental evidence that thermal selection shapes mitochondrial genome evolution

**DOI:** 10.1101/133389

**Authors:** Zdeněk Lajbner, Reuven Pnini, M. Florencia Camus, Jonathan Miller, Damian K. Dowling

**Author notes:** Correspondence: Zdeněk Lajbner, Physics and Biology Unit, Okinawa Institute of Science and Technology Graduate University (OIST), 1919-1 Tancha, Onna-son, Okinawa 904-0945, Japan. Competing Interests: The authors declare no competing interests. Data accessibility statement: Data accessible in Supplementary Table S2.

## Abstract

Mitochondria are essential organelles, found within eukaryotic cells, which contain their own DNA. Mitochondrial DNA (mtDNA) has traditionally been used in population genetic and biogeographic studies as a maternally-inherited and evolutionary-neutral genetic marker. However, it is now clear that polymorphisms within the mtDNA sequence are routinely non-neutral, and furthermore several studies have suggested that such mtDNA polymorphisms are also sensitive to thermal selection. These observations led to the formulation of the “mitochondrial climatic adaptation” hypothesis, for which all published evidence to date is correlational. Here, we use laboratory-based experimental evolution in the fruit fly, *Drosophila melanogaster,* to test whether thermal selection can shift population frequencies of two mtDNA haplogroups whose natural frequencies exhibit clinal associations with latitude along the Australian east-coast. We present experimental evidence that the thermal regime in which the laboratory populations were maintained, drove changes in haplogroup frequencies across generations. Our results strengthen the emerging view that intra-specific mtDNA variants are sensitive to selection, and suggest spatial distributions of mtDNA variants in natural populations of metazoans might reflect adaptation to climatic environments rather than within-population coalescence and diffusion of selectively-neutral haplotypes across populations.

**Impact Summary:** We applied experimental laboratory evolution to provide the first direct test of the “mitochondrial climatic hypothesis,” which predicts that the variation of mitochondrial genomes across natural distributions of metazoans can be shaped by thermal selection. Our design is the first of its kind when it comes to inferring the role of thermal selection in shaping mtDNA frequencies in nature. We harness two naturally occurring mtDNA haplotypes of *Drosophila melanogaster* that segregate along the east coast of Australia. One of these haplotypes predominates at sub-tropical northern latitudes and the other in the temperate and cooler south of the country. We then compete these haplotypes against each other in replicated experimental fly populations submitted to one of four different thermal regimes, in either the presence or absence of infection by *Wolbachia*, a coevolved endosymbiont that also exhibits maternal transmission.

We confirm that when evolving in the laboratory under warmer conditions, a haplotype naturally predominating in subtropical conditions outcompetes a haplotype that predominates at cooler Australian latitudes in the wild. We see this effect on haplotype frequencies in females in populations where latent *Wolbachia* infections had been purged.

Our results also suggest that sex-specificity of mtDNA effects, and co-occurrence of other maternally-inherited microbiotic entities - of which *Wolbachia* is just one example - are likely to shape the trajectories of mitochondrial genome evolution in the wild.

## Introduction

Mitochondrial DNA (mtDNA) is usually maternally inherited [1], and was long considered a neutral evolutionary marker [2]. Accordingly, the mtDNA has been routinely harnessed as a quintessential tool in phylogenetics, population genetic studies, and especially in phylogeographic reconstructions seeking to understand demographic responses to postglacial climate change [3]. For example, in 2017 alone, more than 1500 studies were published that relied at least in part on mtDNA for phylogenetic and phylogeographic inference, based on the ISI *Web of Science* Core Collection search: mitochondrial AND (phylogeography OR phylogeny). Nevertheless, non-neutral evolution of DNA can compromise historical inferences in population and evolutionary biology [4]. Selection on standing genetic variation plays an important role in facilitating rapid adaptation to novel environments [5, 6]. New evidence published over the past two decades has suggested that a sizeable amount of genetic variation within the mitochondrial genome is sensitive to natural selection, and exerts strong effects on the phenotype [7,8,9,10,11]. Furthermore, emerging data indicate that not all mitochondrial haplotypes perform equally well under the same thermal conditions – some perform best when it is warmer, others when it is colder [12,13,14,15,16]. Correlative molecular data in humans are also consistent with the idea that certain mitochondrial mutations might represent adaptations to cold climates [17,18,19,20], and thus support is growing for a “mitochondrial climatic adaptation” hypothesis, which suggests that polymorphisms that accumulate across mtDNA haplotypes found in different spatial locations have been shaped by selection to the prevailing climate.

These ideas remain contentious, primarily because the conclusions of previous studies are based on correlations between mutational patterns in the mtDNA sequence and climatic regions, which have proven difficult to replicate in other or larger datasets [21, 22]. We therefore decided to apply experimental evolution to test the mitochondrial climatic adaptation hypothesis by determining whether multigenerational exposure of replicated populations of fruit flies to different thermal conditions leads to consistent changes in the population frequencies of naturally-occurring mtDNA haplotypes.

In the wild, different locally-adapted populations routinely come into secondary contact and hybridize, which enables selection of novel mito-nuclear genotypes that might be better suited to a new or changing environment [23]. Such an evolutionary scenario is likely to have become increasingly common in the Anthropocene, wherein humans have rapidly altered both climatic conditions and levels of habitat connectivity [24]. We reproduced such a hybridization event under controlled laboratory conditions by interbreeding two subpopulations of *D. melanogaster,* each adapted to thermal environments at a different end of an established and well-studied latitudinal cline [25, 26]. It is thought that the species was introduced into Australia during the past one to two hundred years, probably via recurrent introductions of flies from both African and European origins [26, 27]. The species has been studied extensively in the context of thermal adaptation along latitudinal clines, both within Australia, and other replicated clines in other continents [25,26,28]. This research has shown that numerous phenotypes related to thermal tolerance exhibit linear associations with latitude, and that these patterns are underscored by linear associations of key candidate nuclear genes [25]. Yet, no research had focused on the quantitative spatial distribution of mtDNA variants [28], until Camus et al. [29] reported that similar clinal patterns are found for two phylogenetic groups of mtDNA haplotypes (haplogroups) along the eastern coast of Australia. Furthermore, Camus et al. [29] were able to map these clinal patterns of mtDNA variation to the phenotype, showing that the mtDNA haplotype that predominates at subtropical latitudes confers superior resistance to extreme heat exposure, but inferior resistance to cold exposure than its temperate-predominant counterparts.

To characterize our model system in detail, we designed a study based on experimental evolution, in which we submitted replicated laboratory populations of *D. melanogaster* to one of four different regimes of thermal selection. We note that similarly to patterns observed in mtDNA haplotype frequencies, *Wolbachia* infection frequencies also concord to latitudinal clinal patterns along the Australian east coast distribution of *D. melanogaster*, with higher frequencies in low latitude populations [30]. Furthermore, both the mtDNA and *Wolbachia* are maternally-inherited. Therefore, it is possible that previously reported clinal patterns in mtDNA haplotypes in Australia [29] might have been in part shaped by direct selection on *Wolbachia* genomes, with changes in mtDNA haplotype frequencies brought about by genetic hitchhiking on particular strains of *Wolbachia*. In order to test the interacting effect of *Wolbachia* infection on the dynamics of mtDNA adaptation under thermal selection, we replicated our experiment, under two different conditions – one in which the ancestors of our experimental flies had been treated with antibiotics to remove *Wolbachia* infections, and the other in which the ancestors had not received antibiotic treatment.

## Material and Methods

### Experimental procedures

Wild subpopulations of *D. melanogaster* were sampled during January 2012 in Australia. We sampled a “hot” adapted subpopulation (“H”; Townsville: −19.26, 146.79) in the northeast, and a “cool” adapted subpopulation (“C”; Melbourne: −37.99, 145.27) in the south of the continent. We collected fertilised females and established 20 isofemale lineages from each wild population. Each lineage then underwent three generations of acclimatisation to laboratory conditions.

Wild fruit flies are often hosts of intracellular parasites, such as *Wolbachia* and associated maternally-transmitted microbiomes that are known to manipulate host phenotypes and affect their thermal sensitivity [31,32,33]. To assess the effects of thermal selection on the standing mitochondrial variation in our experiment, both in the presence and especially the absence of these maternally-inherited microbiota that co-transmit with the mtDNA, we treated a full copy of our isofemale lineages with the antibiotic tetracycline hydrochloride (0.164 mg mL^-1^ tetracycline in food for 3 generations), such that we maintained a one full copy with putative *Wolbachia* and unperturbed microbiomes, and one full copy without *Wolbachia* but with perturbed microbiomes [34].

We then propagated these lineages for a further 10 generations to mitigate any effects of the antibiotic treatment under laboratory conditions. Flies were reared at 25°C on a 12:12 hour light:dark cycle in 10 dram plastic vials on a potato-dextrose-agar medium, with live yeast added to each vial *ad libitum*. All isofemale lineages were then transferred from our laboratories in Australia to those in Japan, and their food medium changed to a corn flour-glucose-agar medium (see Supplementary Table S1), with live yeast added to each vial *ad libitum*. To set up a series of replicated experimental populations, they acclimatized for a further 3 generations at 25°C before entering the admixture process described below.

We pooled 5 virgin females (♀) from each of 18 of the H isofemale lineages, mentioned above, with 5 virgin males (♂) from each of 18 of the C lineages in one bottle (HC = 18 x 5 ♀H + 18 x 5 ♂C), and 5 virgin females from each of the 18 C isofemale lineages with 5 virgin males from each of the 18 H isofemale lineages in another bottle (CH = 18 x 5 ♀C + 18 x 5 ♂H), such that each bottle contained 90 males and 90 females. This step was performed separately for flies sourced from the tetracycline-treated isofemale lineages and flies sourced from untreated isofemale lineages (Fig. 1; Supplementary Table S2). We then allowed the flies to lay eggs over 8 consecutive days and transferred them to fresh bottles as indicated in Supplementary Table S3. From this step, we reared all flies in 250 ml bottles on a corn flour-glucose-agar medium until the experiment was concluded. In the next step, we created each experimental population by mixing F1 offspring (25 virgin males and 25 virgin females) from the HC bottles with corresponding F1 offspring (25 virgin males and 25 virgin females) from the CH bottles (25 ♂CH + 25 ♀HC + 25 ♀CH + 25 ♂HC). These flies (randomly selected from the first hybrid generation) we call the starting generation. In this way, we established 7 experimental populations from the tetracycline-treated isofemale lineages and 8 experimental populations from the untreated lineages. We allowed flies of these populations to mate and lay eggs at 25°C before transferring bottles with approximately 500 eggs each into four thermal regimes, represented by cold versus warm temperatures on either a constant or fluctuating temperature cycle. This design nests the temperature treatment inside the experimental population; each experimental population consists of four experimental subpopulations, and each experimental subpopulation adapts to one of the four thermal conditions. Bottles maintained in the constant cold temperature were kept at 19°C (denoted “19°C”), and those in the constant warm temperature at 25°C (denoted “25°C”). In addition, we used Environmental Chambers (MIR-154, Sanyo) to generate fluctuating thermal conditions that are common in areas of origin of our experimental populations [35]. The first such conditions (denoted “cold”) are typical of Melbourne (mean of 17.4°C): 8:00(22°C); 11:30(28°C); 16:00(20°C); 20:00(17°C); 22:00(14°C); 8:00(15°C); 11:30(20°C); 16:00(16°C); 20:00(15°C); 22:00(14°C). The second (denoted “warm”) are typical of Townsville (mean of 26.4°C): 8:00(27°C); 10:30(28°C); 20:00(27°C); 22:30(26°C); 0:00(24°C); 8:00(26°C); 10:30(28°C); 20:00(27°C); 22:30(26°C); 0:00(25°C). The temperatures in all conditions were continually monitored, and in fluctuating conditions recorded by Thermo-hydro SD Data Loggers (AD-5696; A&D Ltd). We propagated all replicate populations for three months (3 or 7 successive discrete generations depending on the thermal condition; Supplementary Table S4). We controlled the size of each subpopulation by trimming egg numbers in each generation to approximately 500. At the end of the experimental evolution period of three months, adult flies were collected and fixed in 95% ethanol.

**Figure 1:**
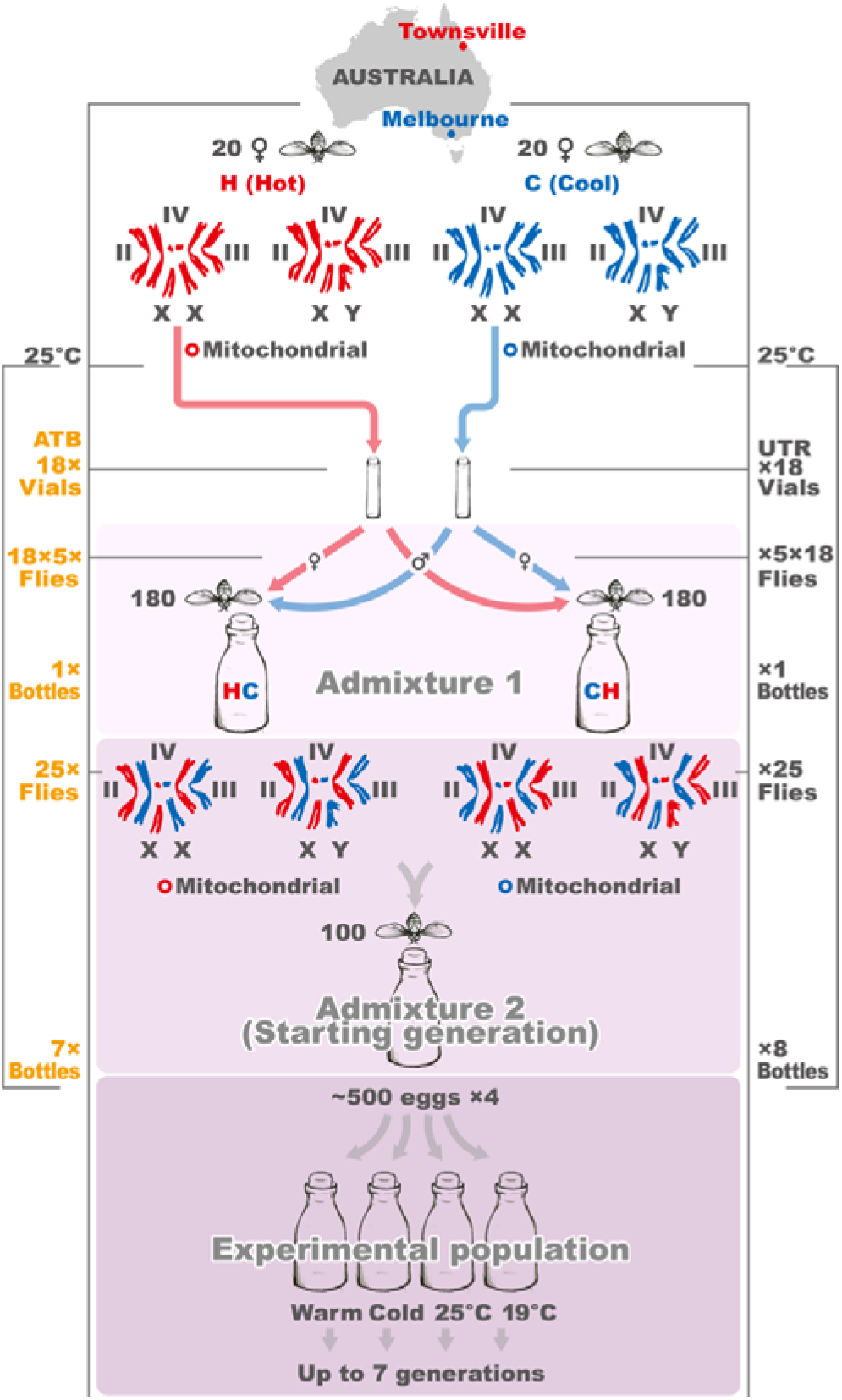
Scheme of experimental evolution by hybridization of differentially thermally-adapted subpopulations of fruit fly. Prior to the application of thermal selection, we created a series of replicated experimental populations, by combining flies of isofemale lineages collected from the Melbourne (putatively cool-adapted, or “C”) subpopulation, denoted in blue, and the Townsville (putatively hot-adapted, “H”) subpopulation (red). This was achieved over two generations, via a process of admixture of the individual isofemale lineages. In the Admixture 1 step, we pooled 5 virgin females (♀) from each of 18 of the H isofemale lineages, with 5 virgin males (♂) from each of 18 C isofemale lineages into one bottle, denoted by HC = 18 x 5(♀H) + 18 x 5(♂C). In parallel in Admixture 1, we performed the reciprocal cross wherein H <=> C above, denoted by CH = 18 x 5(♀C) + 18 x 5(♂H). Each bottle contained 90 males and 90 females (180 flies). In the following generation, at Admixture step 2, we combined 25 virgin females and 25 virgin males from HC bottles together with 25 virgin females and 25 virgin males from CH bottles, 25 (♂CH) + 25(♀HC) + 25(♀CH) + 25(♂HC), across 15 biological replicates (7 of which were descendants of flies treated by antibiotics, 8 of which were descendants of untreated flies). At this stage, all flies had been maintained in standard laboratory conditions (25°C) for 16 generations (14 generations as isofemale lineages, 2 during the admixture process). We then divided each of these 15 biological replicates into 4 subpopulations, subjecting each subpopulation to one of four thermal treatments (19°C, 25°C, fluctuating cold, and fluctuating warm), with each experimental subpopulation containing around 500 individuals. On the left side of the figure, yellow text denotes sample sizes associated with each stage of the admixture process for flies whose ancestors had been exposed to antibiotic treatment (ATB), while grey text on the right corresponds with untreated flies (UTR).

### Data collection

Total genomic DNA was extracted using DNeasy Blood & Tissue Kit (Qiagen). We sequenced total DNA of H and C population samples quantitatively, using an Illumina platform at Micromon (Monash University, Australia). Length of reads was set to 70bp and we reached a maximum coverage 500x on coding parts of mitogenomes [29].

We mapped all reads to the published mitogenomic sequence NC 001709 in Geneious R6 [36]. We observed overall mitogenomic variability and picked 14 mtDNA polymorphic sites (SNPs) that are not unique to the H or C populations. These SNPs segregate all flies into one of two corresponding mtDNA haplogroups denoted A and B in Camus et al. [29] (Fig. 2, Supplementary Table S5). Using multiplexPCR and MALDI/Tof (mass spectrometry; Geneworks, Australia), we genotyped nearly all flies in the starting generation (50 males and 50 females in each of the 15 experimental populations) and at least 24 males and 25 females per experimental subpopulation in the final generation upon completion of experimental evolution. A lower bound for the number of samples to sequence was estimated assuming α = 0.05 and a relative thermal effect of 10% at a power of 1 – β = 70 %. In total, we genotyped 4410 individuals (data available in Supplementary Table S2).

**Figure 2:**
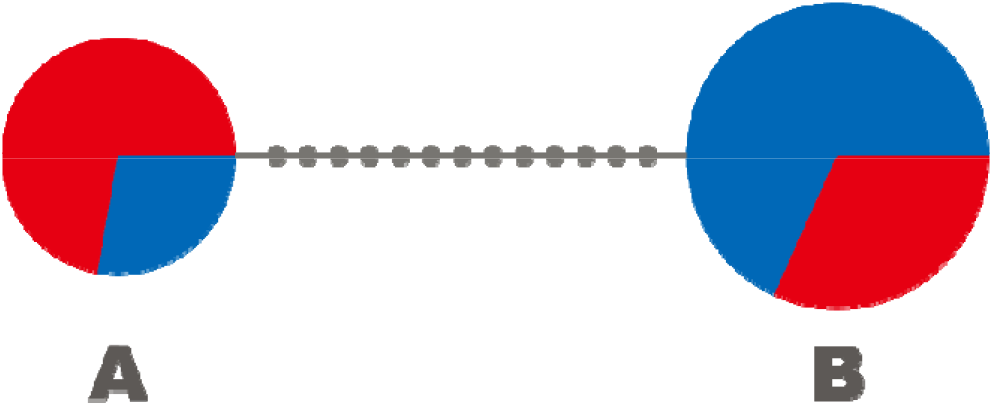
Relationship of A and B mtDNA haplogroups. The circle area for each haplogroup is proportional to its frequency in the wild sample (A=18 females, B=22 females). Colours indicate the sampling region: Townsville (red, 20 females) and Melbourne (blue, 20 females). Small grey circles represent genotyped-SNP divergence (Supplementary Table S5).

### Data analysis

MtDNA is transmitted maternally; males do not transmit their mtDNA to their offspring; and therefore evolutionary changes in mitochondrial genomes must proceed via selection on females. Therefore, our analyses focus on estimating changes in mtDNA haplogroup frequencies in females of each experimental population across the three months of the experiment. We applied a linear mixed-effect model (*lmer*) in the *lme4* v. 1.1-10 package [37] of R version 3.2.2. [38] in RStudio server v. 099.465 [39] with restricted maximum likelihood estimation of variance components, and type III Wald F-tests with Kenward-Roger degrees of freedom appropriate for finite sample size [40]. We modelled the antibiotic and thermal treatment as fixed effects, and the experimental population (n=15 levels) as a random effect. This model revealed a significant interaction between the fixed effects of antibiotic treatment and thermal regime on the frequency change of the B haplogroup (Table 1). To interpret the underlying basis of this interaction, we then decomposed the analysis into two linear mixed-effect models examining haplogroup frequency change according to thermal regime for descendants of flies treated by antibiotics (ATB) and untreated (UTR) separately (Table 2). We modelled the thermal treatment as a fixed effect, and the experimental population as a random effect. Statistical significance between fluctuating-cold and fluctuating-warm thermal treatments has been evaluated by multiple Welch’s t-tests (Table 3). We then estimated selection coefficients according to the haploid selection model and verified that our measurements are not affected by genetic drift by simulations of the Wright-Fisher model (Appendix S1).

**Table 1:** Two-level mixed model comparison of B mtDNA haplogroup frequency change according to antibiotic treatment and thermal regime. Antibiotic treatment, and thermal regime were modelled as fixed effects. Experimental population was modelled as random effect. The green background indicates statistical significance.

|  | *F* | d.f. | P(>*F*) |
| --- | --- | --- | --- |
| antibiotic treatment | 0.0773 | 1 | 0.7821 |
| thermal regime | 0.6730 | 3 | 0.5738 |
| antibiotic treatment: thermal regime | 3.0261 | 3 | 0.0409 |

**Table 2:**
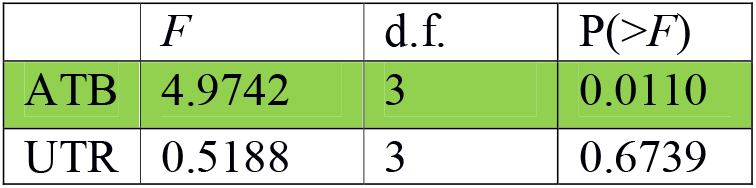
Linear mixed models examining B mtDNA haplogroup frequency change according to thermal regime for descendants of flies treated by antibiotics (ATB) and untreated (UTR) separately. Thermal conditions have been modelled as fixed effects. Experimental population has been modelled as random effect. The green background indicates statistical significance.

**Table 3:** Welch’s t-tests of B mtDNA haplogroup frequency change between thermal regimes for descendants of flies treated by antibiotics (ATB). In the fourth column, the *P*-values are corrected by a Bonferroni factor of 6. The green background indicates statistical significance, especially between fluctuating-cold and fluctuating-warm in accordance with Fig. 3.

|  | t | P(>t) | Modified P |
| --- | --- | --- | --- |
| 19-25 | 1.3176 | 0.1101 | 0.6606 |
| 19-cold | 0.3734 | 0.3580 | 2.1479 |
| 19-warm | 2.9946 | 0.0067 | 0.0404 |
| 25-cold | 1.6366 | 0.0681 | 0.4085 |
| 25-warm | 1.2687 | 0.1154 | 0.6923 |
| cold-warm | 3.4036 | 0.0034 | 0.0202 |

In order to evaluate whether haplogroup frequencies in males are following the frequencies in females in our starting generation, we applied a linear mixed-effect model comparison of haplogroup frequencies between males and females in starting generation according to antibiotic treatment. Antibiotic treatment and sex were modelled as fixed effects; experimental population was modelled as a random effect.

Even within a single generation, thermal selection might yield sex differences in differential survival, from egg to adulthood, of individuals bearing different mtDNA haplotypes; however, under strict maternal inheritance these differences will be reset at each generation. To evaluate the capacity for thermal selection to evoke such within-generation sex-differences in mtDNA frequencies, we applied a multilevel model examining the effect of sex, antibiotic treatment, and thermal regime on B mtDNA haplogroup frequency in the final generation. We modelled the thermal treatment as a fixed effect, the experimental population (n=15 levels) and sub-populations (n=60 levels) as random effects. Statistical significance of within generation frequency differences between sexes in particular treatments has been evaluated by homoscedastic two-tail t-tests in Microsoft Excel.

## Results

The A haplogroup is found to predominate in the low-latitude, hot, tropical subpopulation from Townsville (H), whilst the B haplogroup predominates in the temperate, cooler Melbourne subpopulation (C; Fig. 2). Starting haplogroup frequencies in our experimental populations reflect the composition of the wild populations. On average, 45% of flies at the outset of the experiment possessed the A haplogroup and 55% the B haplogroup. These frequencies were confirmed by individual genotyping of nearly all flies in all 15 experimental populations, at this starting generation of experimental evolution (Supplementary Table S2).

We observed a statistically significant two-way interaction between thermal regime and antibiotic treatment on changes in haplogroup frequency in our experiment (Table 1). The interaction was driven by an effect of thermal regime on haplotype frequencies in the antibiotic treated, but not the untreated, populations (Group ATB, *P =* 0.0110, Fig. 3, Table 2). This effect is important because in the absence of *Wolbachia* infection, changes in haplogroup frequencies can presumably be attributed directly to selection on standing variation in the mitochondrial genome. In the antibiotic treated group, we found that the frequency of the B haplogroup decreased in both of the warmer treatments but increased in the colder treatments. This response is consistent with the spatial distribution of the haplogroups along the Australian cline, where the B haplogroup predominates at temperate higher latitudes, while the A haplogroup predominates in subtropical low latitudes [29]. The largest statistically significant haplotype frequency differences are observed between fluctuating cold and fluctuating warm conditions (*P* = 0.0034 in Table 3). We estimated the selection coefficient of the B haplogroup for fluctuating warm conditions s_w_ = −0.082±0.026. We estimated the selection coefficient of the B haplogroup for fluctuating cold conditions s_c_ = 0.085±0.050 (Fig. 4, Appendix S1). Simulations confirm that our observations are unlikely to be accounted for solely by drift (*P*= 0.0013 in Appendix S1).

**Figure 3:**
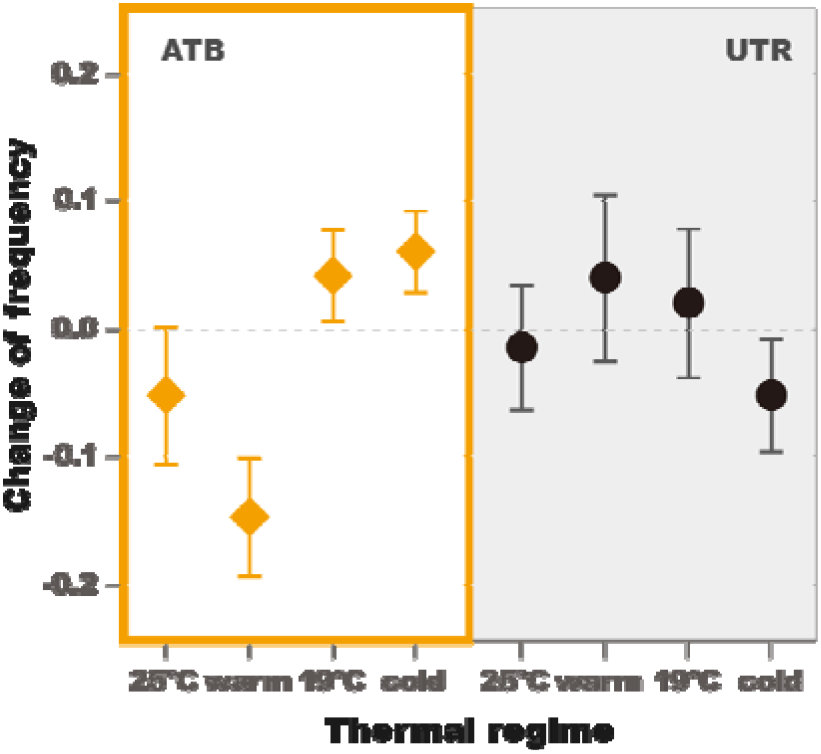
Mean change of B mtDNA haplogroup frequency per thermal environment. Plots depict change in frequencies (final generation-initial generation) in constant **25°C**, fluctuating **warm,** constant **19°C**, and fluctuating **cold** environments for female descendants of flies treated by antibiotics (ATB; 7 replicates) and untreated (UTR; 8 replicates; in which *Wolbachia* and associated maternally transmitted microbiomes present). The error-bars are estimated as ^—^where is the sample standard deviation and N the number of samples.

**Figure 4:**
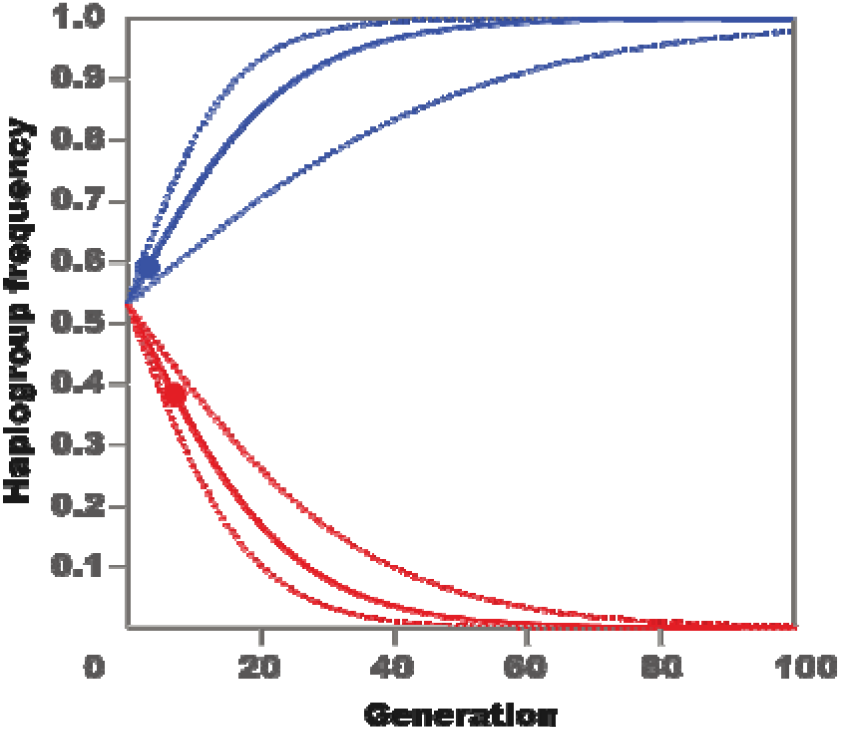
Estimated change in B mtDNA haplogroup frequency extrapolated to 100 generations of flies whose ancestors had had their coevolved microbiomes, including *Wolbachia* infection, disrupted by antibiotic treatment (ATB). Colours indicate the experimental fluctuating thermal conditions representing sampling regions: Townsville (red) and Melbourne (blue). Circles mark the actual values we obtained from experiment.

We also observed an effect of the thermal regime on within-generation sex differences in the frequency of the mtDNA haplogroups (Tables 4 and 5, Figs 5 and 6). These differences were already apparent in the starting generation (*P*= 0.0387 in Table 4). The patterns observed in the antibiotic-untreated populations at 25°C at the commencement of the experiment (panel I in Figs 5 and 6) tracked closely those observed at the conclusion of the experiment seven generations later for populations maintained under the same conditions (UTR 25°C, panel II in Figs 5 and 6). This replication of the sex-specificity of mtDNA frequencies is striking, supporting the finding that sex-differences in frequencies are not occurring randomly (*P*= 0.0321 in Table 5). Under these particular conditions, the frequency of the B haplogroup was higher in adult males than in adult females, suggesting differences in egg-to-adult survivorship of the two haplogroups (UTR 25°C, panels I and II in Figs 5 and 6, *P*= 0.0310). Nonetheless, visual inspection of Figs 5 and 6 reveals other instances of sex specificity in haplogroup frequencies across the experimental treatments, including changes in the direction of sex bias across different combinations of thermal regime and antibiotic treatment. For example, in antibiotic-treated populations, frequencies of the B haplogroup exhibited signatures of female-bias under cooler conditions, particularly in the fluctuating cold regime (ATB panel V in Figs 5 and 6, *P*= 0.0058).

**Table 4:** Mixed model comparison of B mtDNA haplogroup frequencies according to antibiotic treatment between males and females in the starting generation. Antibiotic treatment, and sex were modelled as fixed effects. Population was modelled as a random effect. The green background indicates statistical significance.

|  | *F* | d.f. | P(>*F*) |
| --- | --- | --- | --- |
| sex | 5.5410 | 1 | 0.0337 |
| antibiotic treatment | 0.1889 | 1 | 0.6683 |
| sex: antibiotic treatment | 5.2061 | 1 | 0.0387 |

**Table 5:**
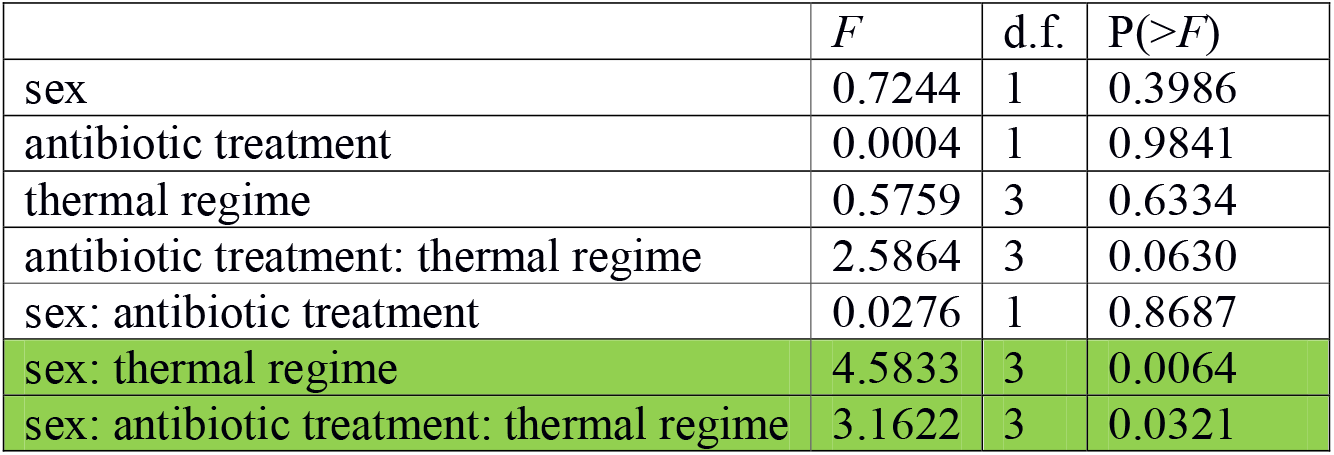
Multilevel model examining the effect of sex, antibiotic treatment, and thermal regime on B mtDNA haplogroup frequency in final generation, as a response variable. Sex, antibiotic treatment, and thermal regime were modelled as fixed effects. Experimental subpopulation and Experimental population were modelled as random effects. The green background indicates statistical significance.

|  | *F* | d.f. | P(>*F*) |
| --- | --- | --- | --- |
| sex | 0.7244 | 1 | 0.3986 |
| antibiotic treatment | 0.0004 | 1 | 0.9841 |
| thermal regime | 0.5759 | 3 | 0.6334 |
| antibiotic treatment: thermal regime | 2.5864 | 3 | 0.0630 |
| sex: antibiotic treatment | 0.0276 | 1 | 0.8687 |
| sex: thermal regime | 4.5833 | 3 | 0.0064 |
| sex: antibiotic treatment: thermal regime | 3.1622 | 3 | 0.0321 |

**Figure 5:**
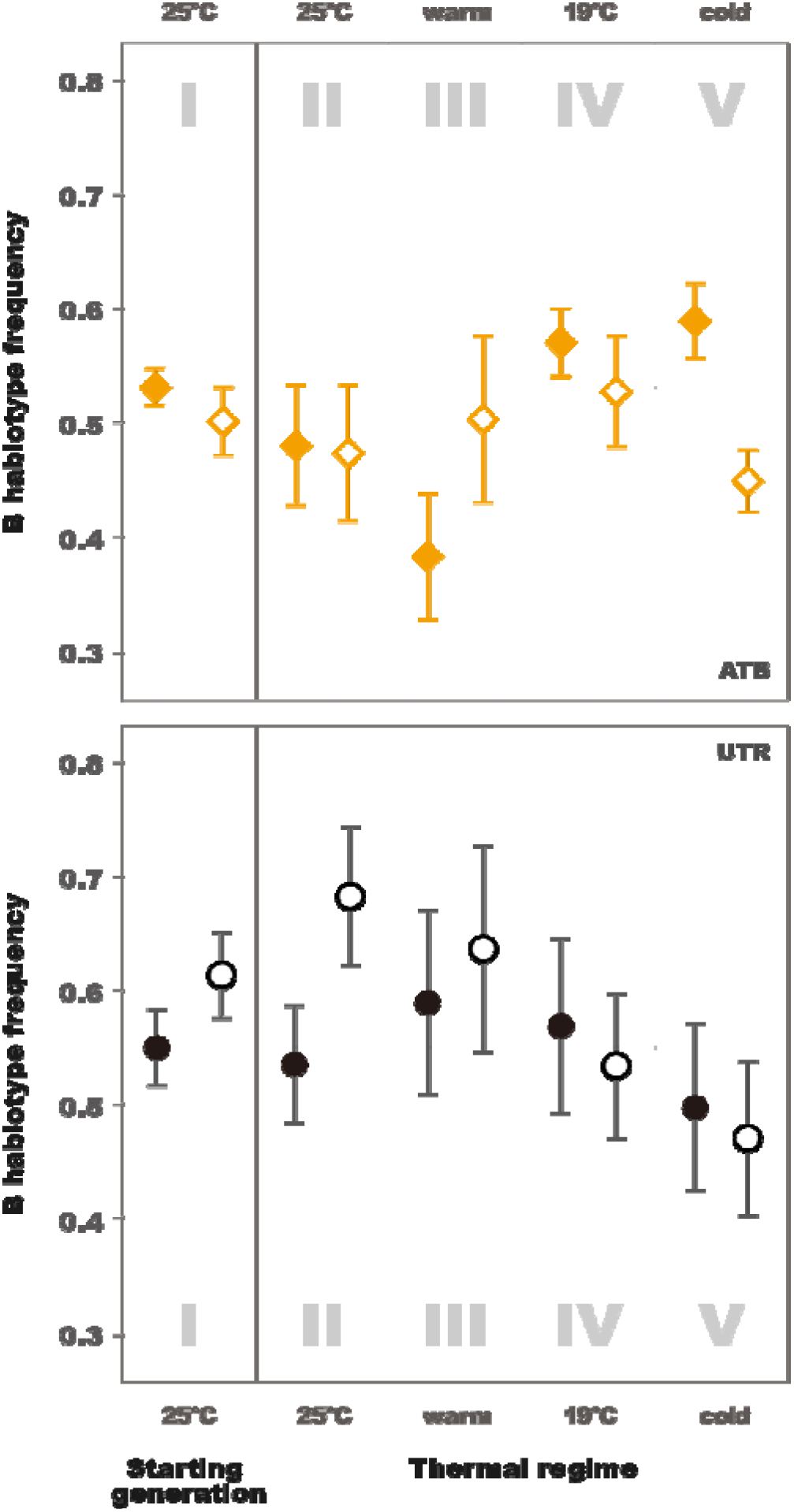
B mtDNA haplogroup frequency per thermal environment. Plots depict frequencies in starting generation (I) and final generation constant **25°C** (II), fluctuating **warm** (III), constant **19°C** (IV), and fluctuating **cold** (V) environments for female (filled shape) and male (empty shape) descendants of flies treated by antibiotics (ATB; yellow squares, 7 replicates) and untreated (UTR; black circles, 8 replicates; in which *Wolbachia* and associated maternally transmitted microbiomes present). Dashed line marks mean starting B mtDNA haplogroup frequency in females. The error-bars are estimated as ^—^where is the sample standard deviation and N the number of samples.

**Figure 6:**
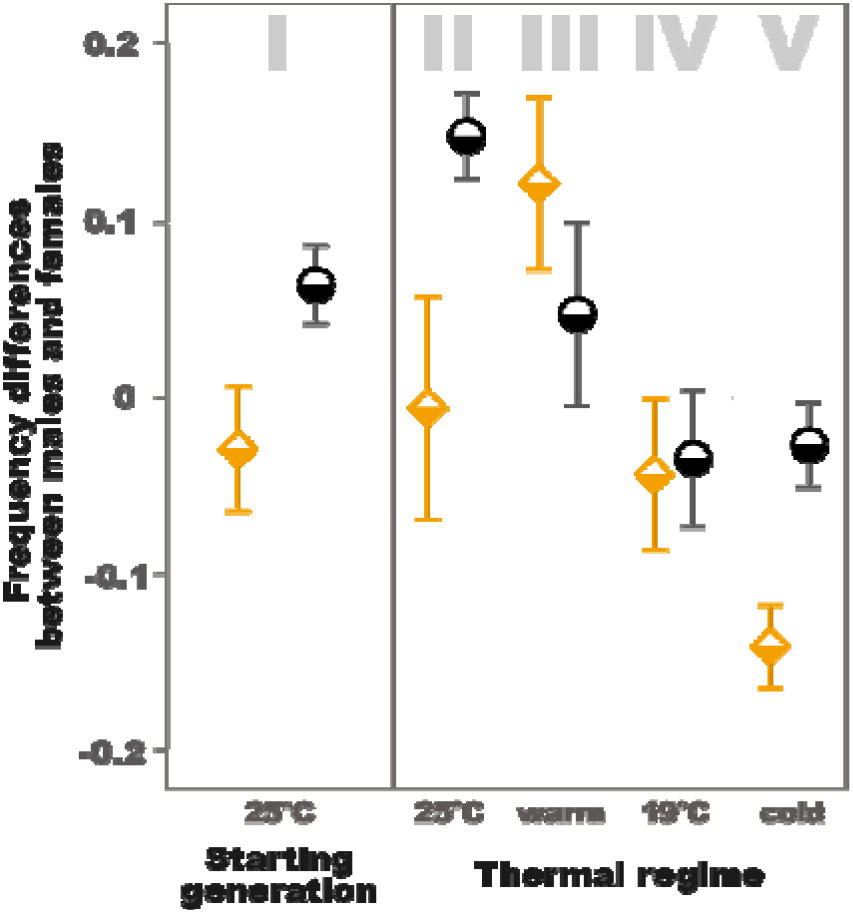
Mean differences of B mtDNA haplogroup frequency between sexes per thermal environment. Plots depict frequencies differences (within generation males - within generation females) in starting generation (I) and final generation constant **25°C** (II), fluctuating **warm** (III), constant **19°C** (IV), and fluctuating **cold** (V) environments for descendants of flies treated by antibiotics (ATB; yellow squares, 7 replicates) and untreated (UTR; black circles, 8 replicates; in which *Wolbachia* and associated maternally transmitted microbiomes present). The error-bars are estimated as ^—^where is the sample standard deviation and N the number of samples.

## Discussion

Our experimental evidence demonstrated context-dependent shifts in population frequencies of two naturally-occurring mtDNA haplogroups under thermal selection. This result is broadly consistent with the hypothesis that spatial distributions of mtDNA haplotypes in natural populations might be in part shaped by thermal selection. In experimental populations whose ancestors’ coevolved microbiomes, including *Wolbachia* infection, were disrupted by antibiotic treatment (ATB), we observed significant effects of thermal selection on mtDNA haplogroup frequencies, in patterns consistent with their corresponding frequencies in the wild. The advantage of these conditions (ATB) is that we could attribute frequency changes of mtDNA haplogroups directly to the variation in the thermal selection regime. Furthermore, our estimated selection coefficients would predict that in the absence of demographic or genetic processes other than thermal selection in shaping the haplogroup distributions (Fig. 4), the mtDNA haplogroups would reach reciprocal mitochondrial fixation in less than 10 years on both sides of the eastern Australian thermal cline.

Nevertheless, haplogroup frequencies have not reached fixation in wild populations along the Australian latitudinal cline [29]. Indeed, twenty years prior to the study of Camus et al. [29], Boussy et al. [41] reported that both the A and B haplogroups coexisted within most sampled populations along the Australian eastern seaboard. Haplogroup A possesses the *HinfI* restriction site used in their study (see Supplementary Table S5). Clearly the dynamics of selection in the wild will differ from those in a highly controlled laboratory experiment, and spatial and temporal environmental variation is likely to lead to genotype-by-environment interactions that might maintain these mitochondrial haplotypes in the wild notwithstanding their sensitivity to thermal selection. In agreement with this argument, our experiments found that the predicted patterns of the A haplogroup outcompeting the B haplogroup under warmer conditions were sustained only in experimental populations that had been antibiotic-treated, and had therefore experienced a microbiome perturbation, and were free from *Wolbachia*. We did not observe the predicted patterns in populations in which flies harboured their coevolved microsymbionts, including possible infections with *Wolbachia*, which might have for instance lead to *Wolbachia*-induced cytonuclear incompatibilities [42, 43] and diverse fitness related effects [31, 44]. *Wolbachia* infection transmission dynamics are known to depend on thermal conditions [45], and thus thermal selection in these untreated populations would presumably be mediated through complex interactions between host nuclear genome, mtDNA haplotype, *Wolbachia* genome, microbiont genomes, which might have obscured any direct effects on the mtDNA frequencies. Furthermore, in the wild, there is a latitudinal cline in *Wolbachia* presence [44], indicating that *Wolbachia* prevalence is likely to be itself shaped by climatic selection. The low-latitude Australian sub-tropical populations exhibit higher levels of *Wolbachia* infection than higher latitude temperate populations [30, 44]. It is possible that genetic polymorphisms within the mitochondrial genome will interact with polymorphisms spanning distinct *Wolbachia* strains to affect fitness outcomes of their hosts. The nature of these complex interactions between *Wolbachia*, mitochondrial, and host nuclear genome currently remains completely unexplored. In the absence of further research that elucidates the relative contributions of selection on *Wolbachia* versus mtDNA haplotypes in shaping the patterns of mitochondrial haplotypic variation observed in nature, it remains difficult to derive predictions as to how host-*Wolbachia* dynamics will affect changes in mito-genomic compositions of natural fruit fly populations [30, 46]. *Wolbachia* clades are also known to exhibit habitat-specific fitness dynamics [47], and it is possible that different *Wolbachia*, or other microsymbiont, strains are linked to the two different mtDNA haplogroups studied here, given that each co-transmit with the mtDNA in perfect association along the maternal lineage, and that the mtDNA frequencies in the antibiotic-free treatments hitchhiked on frequency changes involving these microsymbiotic assemblages, as is expected by theory, and has been observed previously [48, 49].

We observed that haplotype frequencies in males did not necessarily track frequencies in females across the experimental treatments. These sex differences across the experimental treatments were complex, and involved changes in sign under the different combinations of experimental conditions. A key point to note is that selection on mtDNA in males will not directly contribute to shaping patterns of mtDNA variation between generations, under the assumption that males virtually never transmit their mtDNA haplotypes to their offspring. As such, mitochondrial genomes are predicted to evolve under a sex-specific selective sieve [50], in which mutations in the mtDNA sequence that confer harm to males can nonetheless accumulate in wild populations, so long as these same mutations are neutral or beneficial for females [51,52,53,54]. In the absence of inter-sexual positive pleiotropy, such male-expression specific mtDNA mutations could in theory shape patterns of haplotype frequencies within a generation, if they affect male-specific patterns of juvenile or adult survival, but would not be passed on to the next generation, and would thus not shape haplotype frequencies across generations. Male-biased mitochondrial genetic effects on key life history phenotypes have been observed previously [50,54,55,56,57,58], and the patterns observed here strengthen the emerging view of ubiquity of sex-specific effects of mtDNA polymorphisms and suggest that such polymorphisms are sensitive to selection imposed by the thermal and microbial environment.

It is possible that paternal leakage (transmission of mtDNA haplotypes from fathers to offspring) could have affected the evolutionary dynamics of selection across the experimental treatments. While paternal leakage has been previously observed in *Drosophila*, most reports have been limited to cases involving interspecific crosses between individuals of divergent species [59, 60], or intraspecific crosses between individuals with high levels of genetic divergence [61]. One recent study, which sampled flies from natural European and Mediterranean populations of *D. melanogaster*, noted that as many as 14% of individuals were heteroplasmic for divergent haplotypes, thus indicating paternal leakage [62]. Notwithstanding, the average frequency of the minor haplotype was very low within individuals (less than 1%), and such a low level of heteroplasmy seems unlikely to be of evolutionary significance. Indeed, the Dowling lab had not observed even a single case of paternal leakage among the numerous mitochondrial strains created or maintained in the laboratory over the past decade, despite continual backcrossing to males of isogenic strains possessing a different mtDNA haplotype [29,54,63]. However, in 2017, Wolff et al. [64] observed heteroplasmic individuals in two of 168 replicate populations (1%) following a large experimental evolution study in which flies of two divergent mtDNA haplotypes coexisted across 10 generations. In summary, while paternal leakage appears to occur in this species at low frequencies, we assume at this stage that selection on males would have played at most only a minor role in shaping the intergenerational changes in the frequencies of each haplotype.

Mitochondrial genetic markers remain an important tool for population genetics, despite growing experimental evidence that mitochondrial genetic variation is affected by thermal [29], and other kinds of selection [65]. The evolutionary trajectories of distinct mitochondrial haplotypes might furthermore be selected together with functionally-linked nuclear gene complexes [66, 67]. This reinforces the point that phylogenetic, population-genetic, and biogeographic studies involving mtDNA should incorporate statistical tests to investigate the forces shaping sequence variation and evolution [68], and examine variation at multiple genetic loci [69]. To date, researchers have focused mainly on the effects of nonsynonymous mutations in the evolutionary dynamics of mitochondrial genomes [70], but growing evidence suggests that mitochondrial molecular function is also affected by single nucleotides in synonymous and non-protein coding positions on mtDNA [29]; a contention supported by the current study since there are no non-synonymous SNPs separating the A and B haplogroups [29].

Our study advances understanding of the dynamics of evolutionary adaptation by providing experimental evidence that thermal selection can act upon standing variation in the mtDNA sequence. However, further research is needed to resolve the dynamics of this thermal evolution; for instance, by determining whether thermal selection acts on the mtDNA sequence directly, or on epistatic combinations of mitochondrial-nuclear genotype or mitochondrial-microbial genomes; and whether thermal selection is the main driver of adaptive variation that we see within the mitochondrial genome or whether other environmental variables, such as the nutritional environment [71], are key. Furthermore, it remains unclear how much of the pool of non-neutral genetic variation that delineates distinct mitochondrial haplotypes has actually been shaped by adaptive relative to non-adaptive processes. Finally, because of the difficulty of implementing experimental evolution in vertebrates, almost all experimental work investigating the adaptive capacity of the mitochondrial genome has been conducted on a small number of model invertebrate species [16,53,65,72,73,74,75], with few exceptions [76,77,78]. Future studies should involve a combination of ecological and experimental evolutionary approaches, with high resolution transcriptomics and proteomics applied more generally across eukaryotes, and the development of tests enabling us to reliably discern the footprint of thermal selection in wild populations [79].

## Acknowledgements

We thank Vanessa Kellerman and Winston Yee for assistance with wild sample collection, and Mary Ann Price, Carla Sgrò, Ritsuko Suyama, Garth Illsley, Richard Lee, Nicholas Luscombe, Pavel Munclinger, Takeshi Noda, and Oleg Simakov for helpful advice. We thank Yuan Liu for her assistance with artwork design. This work was supported by the Physics and Biology Unit of the Okinawa Institute of Science and Technology Graduate University (J.M.) and JSPS P12751 + 24 2751 to Z.L. and J.M, the Hermon-Slade Foundation (HSF 15/2) and the Australian Research Council (FT160100022 and DP170100165) to D.K.D. Initial stages of the study were funded by Go8EURFA11 2011003556 to Z.L. and D.K.D.

## Author Contributions

Z.L. and D.K.D. designed the experiment. Z.L. performed the experiment. Z.L. and M.F.C. provided mitogenomic sequences. Z.L., R.P., D.K.D., M.F.C., and J.M. contributed to the data analyses. Z.L., D.K.D., R.P., J.M., and M.F.C. wrote the manuscript.

## Supplementary Information

**Appendix S1: Detailed methods**

### Supplementary Tables Legends

**Supplementary Table S1: Fly food composition**

**Supplementary Table S2: List of samples**

In experimental populations ATB1-ATB7, the ancestors had been exposed to antibiotic treatment, while experimental populations UTR1-UTR8 correspond with untreated flies.

**Supplementary Table S3: Foundation dates of experimental populations.**

Foundation date marks the date at which virgin flies were combined in a bottle as outlined in Admixture Step 2 (Fig. 1) to form the starting generation. In Admixture Step 1, we allowed their parents to lay eggs for about 1 day, and transferred them to a new bottle. This process was repeated across nine days. We call the process by which we transfer the flies to a new bottle a “tip”. Virgin flies of each sex were sourced from several tips, in order to ensure we had an adequate supply of flies to initiate the experimental populations. We show the dates of maternal ovipositioning and virgin collection in a separate column (date). Number means the number of virgin flies sourced from the tip.

**Supplementary Table S4: Propagation of experimental populations in dates.**

Foundation date corresponds with Supplementary Table S3. We imposed the four different thermal regimes on multiple generations starting by eggs laid by the starting generation. Within each generation, the flies of each experimental bottle were transferred to new bottles, and the column “tip number” reflects the “tip” that was used to propagate the next generation per experimental population. Although the tip we used to initiate each experimental population varied in Generation 1, in subsequent generations, we propagated experimental populations mostly from the first tip.

**Supplementary Table S5: Selected SNPs characteristics for mtDNA haplogroups.**

